## Supplementary data for "Structural and biophysical analysis of a *Haemophilus influenzae* tripartite ATP-independent periplasmic (TRAP) transporter"

### Supplementary Tables

**Supplementary Table 1 |** Cryo-EM data collection, processing, refinement, and validation statistics.

| | **Data collection** | |
|---|---|---|
| Instrument | Titan Krios-beta at 300 kV | |
| Detector | K3 BioQuantum | |
| Camera | EF-CCD | |
| Mode | Counted super resolution | |
| Magnification | 130,000x | |
| Pixel size (Å/px) | 0.6645 | |
| C2 aperture (µm) | 50 | |
| Slit width (eV) | 20 | |
| Spot size | 5 | |
| Spherical aberration (mm) | 2.7 | |
| Frames | 163 | |
| Exposure time (s) | 2.18 | |
| Dose | 73.1 | |
| Fractions | 40 | |
| De-focus range | -2.0, -1.8, -1.6, -1.4, -1.2, -1.0, -0.8, -0.6, -0.4 | |
| Shots per hole | 10 | |
| Images collected | 14,281 | |
| Electrons/px/s | 14.798 | |
| Objective aperture | No | |
| | ***Hi*SiaQM (parallel dimer)** | ***Hi*SiaQM (antiparallel dimer)** |
| **Processing** | | |
| Initial particle images (no.) | 2,950,415 | 2,950,415 |
| Final particle images (no.) | 220,810 | 225,044 |
| **Refinement** | | |
| Map resolution (Å) | 3.36 | 2.99 |
| FSC threshold | 0.143 | 0.143 |
| Model composition | | |
| Non-hydrogen atoms | 9521 | 9644 |
| Protein residues | 1224 | 1230 |
| Ligands | 4 | 8 |
| R.m.s. deviations | | |
| Bond lengths (Å) | 0.003 | 0.003 |
| Outliers | 0 | 0 |
| Bond angles (°) | 0.510 | 0.524 |
| Outliers | 1 | 0 |
| **Validation** | | |
| MolProbity score | 1.09 | 1.40 |
| Clash score | 3.02 | 5.25 |
| Rotamer outliers (%) | 0.29 | 0.88 |
| EMRinger score | 1.73 | 3.55 |
| Ramachandran plot | | |
| Favoured (%) | 98.2 | 97.39 |
| Allowed (%) | 1.8 | 2.61 |
| Outliers (%) | 0 | 0 |
| $CC_{volume}$ | 0.80 (0.86*) | 0.86 |
| $CC_{mask}$ | 0.80 (0.86*) | 0.87 |
| $CC_{peaks}$ | 0.60 (0.77*) | 0.70 |
| PDB ID | 8THI | 8THJ |

*values from model fit to consensus parallel dimer map

**Supplementary Table 2 |** Protein sequences of *Hi*SiaQM and *Hi*SiaP expressed and purified in this work.

| **Protein** | **Protein sequence** |
|---|---|
| *Hi*SiaQM<br>https://www.uniprot.org/uniprot/P44543<br>* X denotes the N-terminal purification tag | MGGSHHHHHHGMASMTGGQQMGRDLYDDDDKDRWGSELE<br>MKYINKLEEWLGGALFIAIFGILIAQILSRQVFHSPLIWSEELAKL<br>LFVYVGMLGISVAVRKQEHVFIDFLTNLMPEKIRKFTNTFVQLL<br>VFICIFLFIHFGIRTFNGASFPIDALGGISEKWIFAALPVVAILMM<br>FRFIQAQTLNFKTGKSYLPATFFIISAVILFAILFFAPDWFKVLRIS<br>NYIKLGSSSVYVALLVWLIIMFIGVPVGWSLFIATLLYFSMTRW<br>NVVNAATEKLVYSLDSFPLLAVPFYILTGILMNTGGITERIFNFA<br>KALLGHYTGGMGHVNIGASLLFSGMSGSALADAGGLGQLEIK<br>AMRDAGYDDDICGGITAASCIIGPLVPPSIAMIIYGVIANESIAKL<br>FIAGFIPGVLITLALMAMNYRIAKKRGYPRTPKATREQLCSSFK<br>QSFWAILTPLLIIGGIFSGLFSPTESAIVAAAYSVIIGKFVYKELTL<br>KSLFNSCIEAMAITGVVALMIMTVTFFGDMIAREQVAMRVADV<br>FVAVADSPLTVLIMINALLLFLGMFIDALALQFLVLPMLIPIAMQ<br>FNIDLIFFGVMTTLNMMVGILTPPMGMALFVVARVGNMSVST<br>VTKGVLPFLIPVFVTLVLITIFPQIITFVPNLLIP |
| *Hi*SiaP<br>https://www.uniprot.org/uniprot/P44542<br>* X denotes the periplasmic signal peptide | MMKLTKLFLATAISLGVSSAVLAADYDLKFGMNAGTSSNEYK<br>AAEMFAKEVKEKSQGKIEISLYPSSQLGDDRAMLKQLKDGSLD<br>FTFAESARFQLFYPEAAVFALPYVISNYNVAQKALFDTEFGKDL<br>IKKMDKDLGVTLLSQAYNGTRQTTSNRAINSIADMKGLKLRVP<br>NAATNLAYAKYVGASPTPMAFSEVYLALQTNAVDGQENPLAA<br>VQAQKFYEVQKFLAMTNHILNDQLYLVSNETYKELPEDLQKV<br>VKDAAENAAKYHTKLFVDGEKDLVTFFEKQGVKITHPDLVPF<br>KESMKPYYAEFVKQTGQKGESALKQIEAINP |

### Supplementary Figures

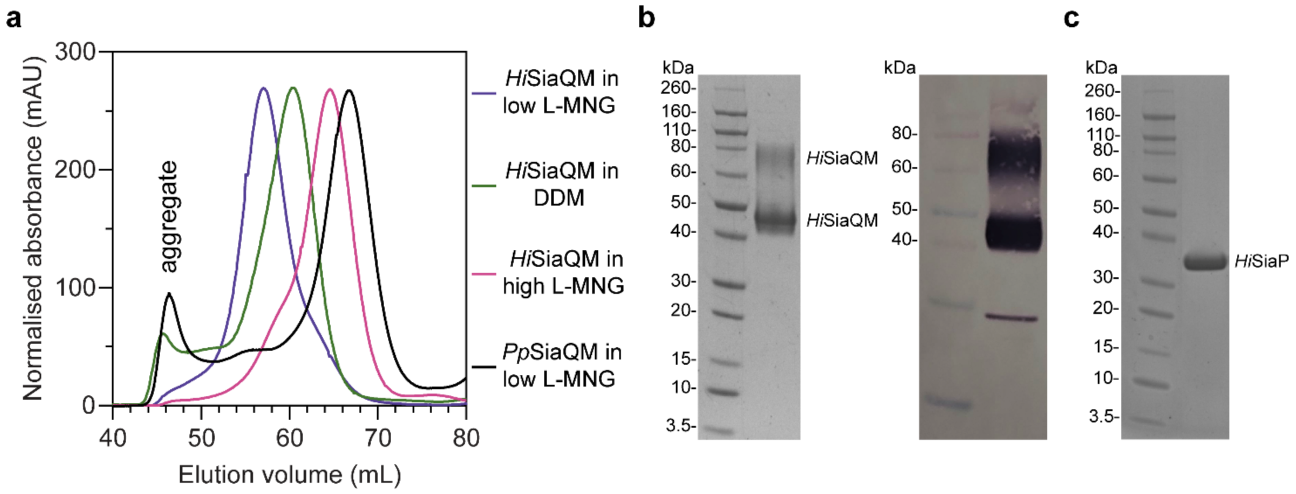


**Supplementary Figure 1 | Size exclusion chromatography traces of SiaQM transporters suggests that *Hi*SiaQM exists as multiple species in detergent.** **a**, *Hi*SiaQM purified in DDM (green trace) results in significant aggregation at 45-50 mL, which is not apparent when solubilised in L-MNG (blue and pink traces). *Hi*SiaQM purified in a low concentration of L-MNG (blue trace) results in a higher order species with a peak at 57 mL compared to *Hi*SiaQM purified in a high concentration of L-MNG (pink trace), which has a peak at 65 mL. At a similar protein concentration (~5 mg/mL) in DDM, *Hi*SiaQM elutes as a single peak between these volumes at 60 mL (green trace). For comparison, non-fused SiaQM from *P. profundum* expressed with the same purification tag and purified in a low concentration of L-MNG (black trace) has a dominant peak at 67 mL, characteristic of a monomer and this greatly contrasts against *Hi*SiaQM in low L-MNG (blue trace). **b**, SDS-PAGE gel (left) of L-MNG purified *Hi*SiaQM that consistently ran as two bands (~45 and ~75 kDa). Protein of this purity (>95%) in L-MNG was used for all subsequent experiments, including for further reconstitution into amphipol or nanodiscs. Western blot (right) with anti-Xpress primary antibody identifies both large bands that appear on SDS-PAGE following purification to be *Hi*SiaQM. A small amount of a lower molecular weight contaminant or degradation product was also present. **c**, SDS-PAGE gel demonstrating the purity of *Hi*SiaP. A single band at ~35 kDa indicates that *Hi*SiaP is greater than 95% pure. Protein of this purity (>95%) was used for all subsequent experiments.

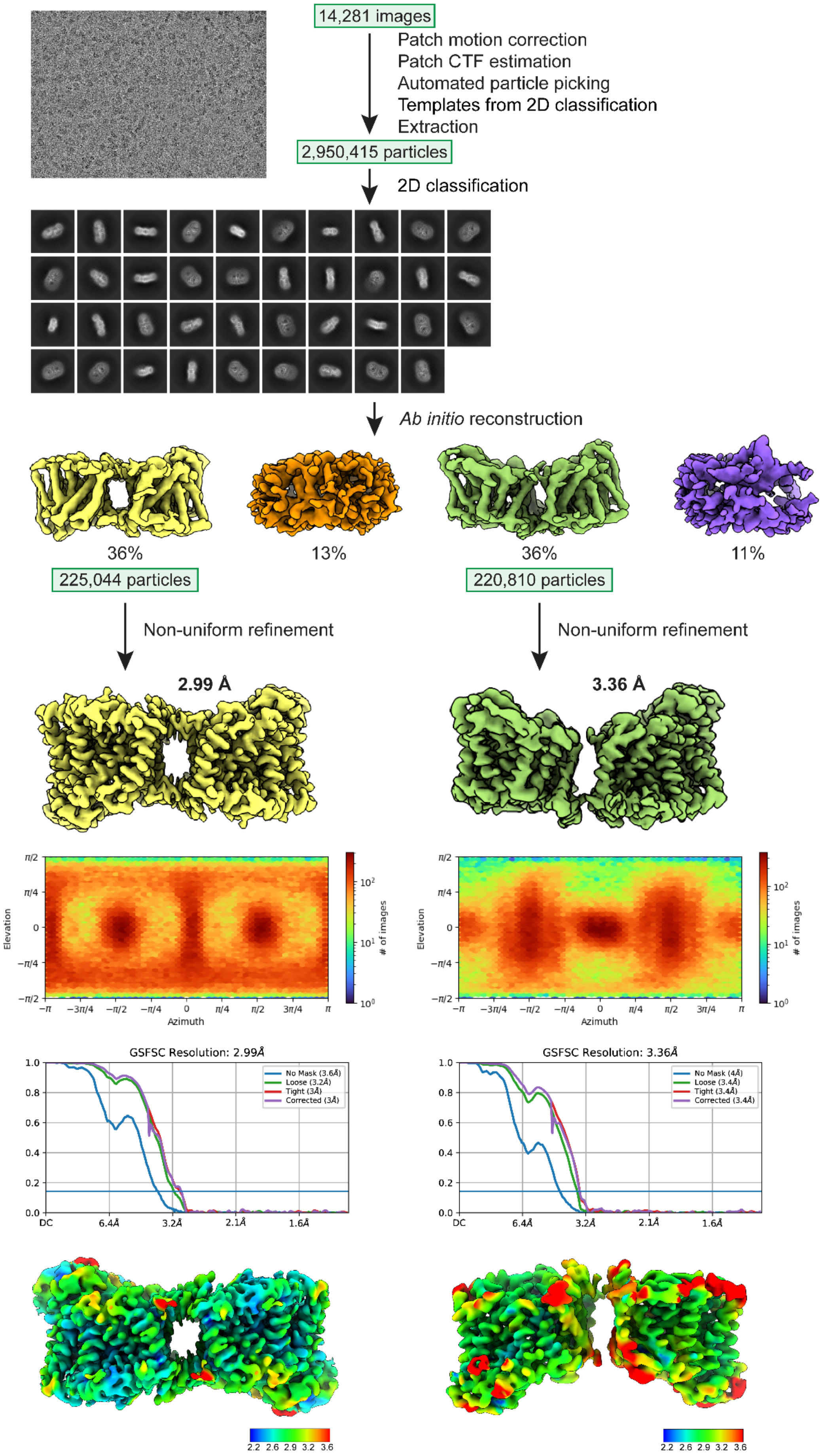


**Supplementary Figure 2 |** Cryo-EM workflow for structure determination. The two classes were determined as 36% of the particles (antiparallel dimer) and 36% (parallel dimer).

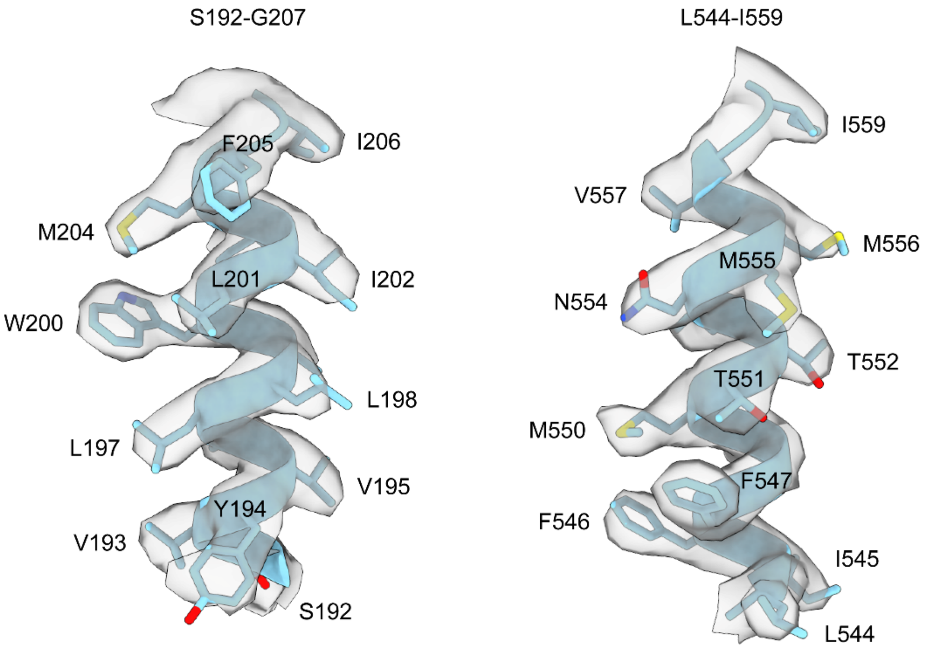


**Supplementary Figure 3 | Representative density of the helices of *Hi*SiaQM.** Clear side chain densities are present for almost all of the residues of *Hi*SiaQM. Density for the antiparallel dimer is shown as it is higher resolution and was used for the structural analysis in this work.

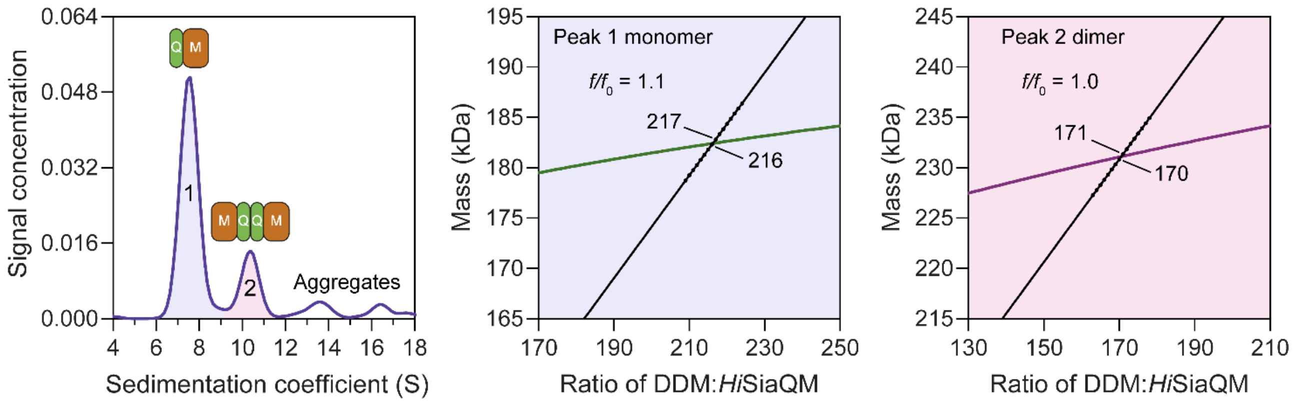


**Supplementary Figure 4 | *Hi*SiaQM self-association in DDM. a**, SV-AUC analysis of *Hi*SiaQM in DDM (left panel). Two well resolved species exist at 7.6 S (diffusion coefficient, D = 5.0 × $10^{-7}$ $cm^2/s$) and 10.3 S (D = 5.0 × $10^{-7}$ $cm^2/s$), with the larger peak constituting 62% of the signal. Due to the presence of protein aggregation in DDM (**Supplementary Figure 1a,** green trace), SV-AUC for this sample was performed at 4 °C (instead of 20 °C for L-MNG) to prevent *Hi*SiaQM from forming larger order species, although even at this temperature larger species are still evident at 12-18 S and extend to ~40 S (not shown). This observed heterogeneity was a significant reason for our use of L-MNG in all other experiments. The species at 7.6 S (Peak 1, blue shading) is most consistent with *Hi*SiaQM as a monomer with ~216 molecules of DDM bound (middle panel; green = measured mass, black = theoretical mass), calculated from the experimental sedimentation and diffusion coefficients. These calculations suggest that Peak 1 existing as a dimer is unlikely, as the dimeric protein would only have ~31 molecules of DDM bound. Additionally, the calculated $f/f_0$ of a monomer for Peak 1 is 1.1, consistent with a protein in a detergent micelle. The species at 10.3 S (Peak 2, pink shading) is most consistent with *Hi*SiaQM as a dimer with ~171 molecules of DDM bound (calculated) (right panel; purple = measured mass, black = theoretical mass); Peak 2 existing as a monomer is not possible, as the protein clearly has a smaller species in Peak 1 and cannot be divided further than a monomer, and a trimer is also unlikely as the experimental data suggests that no DDM would be bound (calculated). Additionally, the calculated $f/f_0$ of a dimer for Peak 2 is 1.0, roughly consistent with a protein in a detergent micelle. These calculations do not account for bound lipid molecules.

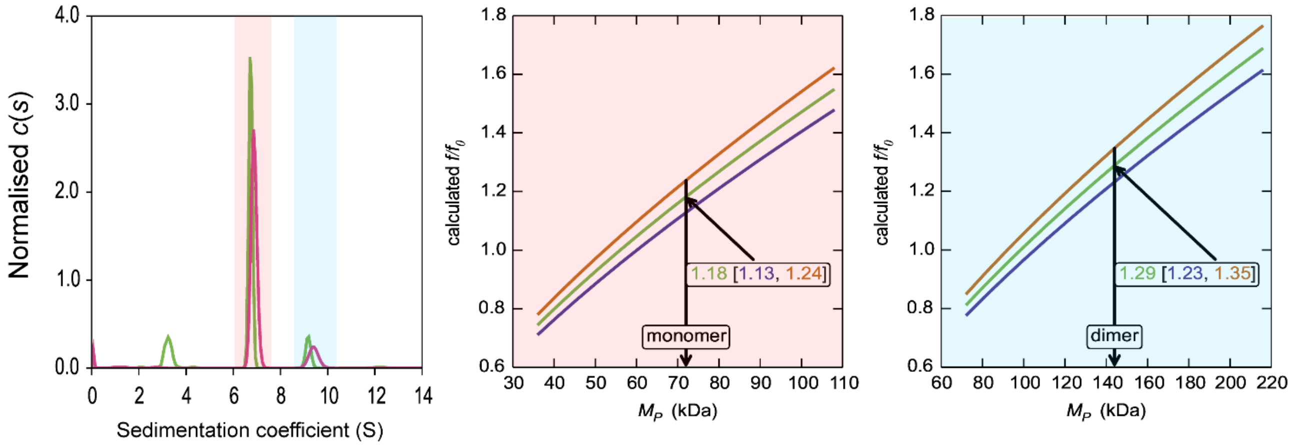


**Supplementary Figure 5 | *Hi*SiaQM self-association in L-MNG (interference analysis).** Sedimentation of *Hi*SiaQM in L-MNG (left panel) with absorbance data (pink) and interference data (green). Two well resolved species exist at 6.7 S and 9.4 S, with the smaller species constituting greater than 80% of the signal. The oligomeric state of the two species was verified using the membrane protein calculations function of GUSSI (77), as described previously (78-81). Free L-MNG micelles are seen only with interference data at ~3.2 S. The peak at 6.7 S is most consistent with a *Hi*SiaQM monomer (middle panel). After determining the amount of L-MNG bound to the protein with the interference data, frictional ratios ($f/f_0$) can be calculated to test the hypothesised oligomeric state. The calculated $f/f_0$ for a monomer for the major species (red shading) is 1.2 (1σ error = 1.13–1.24), consistent with a single *Hi*SiaQM in a detergent micelle. A dimer is unlikely, since the calculated $f/f_0$ for a dimer for the major species is 1.9 (1σ error = 1.79–1.96) (not shown). On the right panel, the calculated $f/f_0$ for a dimer for the minor species (blue shading) is 1.3 (1σ error = 1.23–1.35), again consistent with dimeric *Hi*SiaQM in a detergent micelle. A monomer is unlikely, since the calculated $f/f_0$ for a monomer for the 9.4 S species is 0.8 (1σ error = 0.78–0.85) (not shown). With the detergent bound to the sedimenting protein now known alongside the mass of the detergent monomers, it can be estimated that approximately 90 (monomer) and 150 (dimer) molecules of L-MNG are bound to the protein.

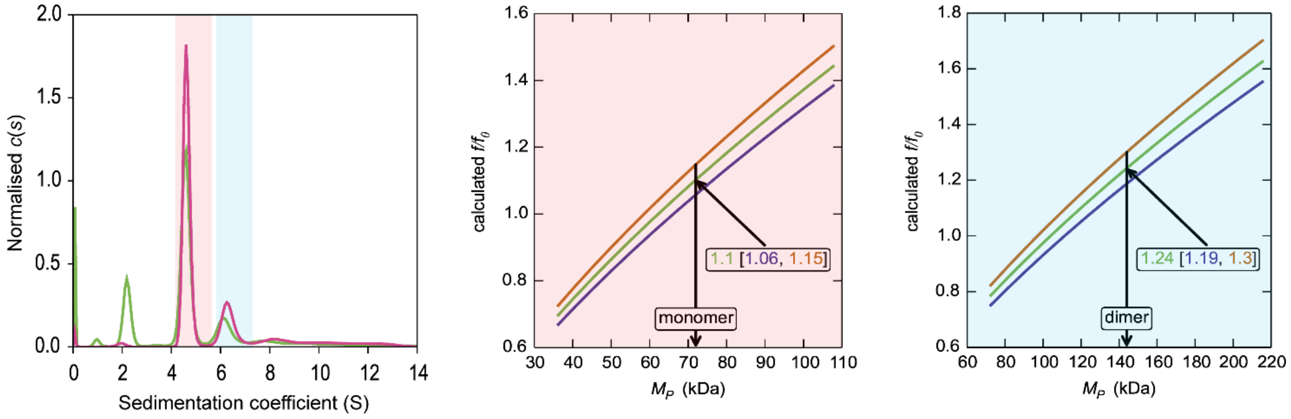


**Supplementary Figure 6 | *Hi*SiaQM self-association in DDM (interference analysis).** Sedimentation of *Hi*SiaQM in DDM (left panel) with absorbance data (pink) and interference data (green). Two main species exist at ~4.5 S and ~6.2 S, with the smaller species constituting greater than 60% of the signal. The oligomeric state of the two species was verified using the membrane protein calculations function of GUSSI (77), as described previously (78-81). Free DDM micelles are seen only with interference data at ~2.1 S. Due to the presence of protein aggregation in DDM (**Supplementary Figure 1a,** green trace), SV-AUC for this sample was performed at 4 °C (instead of 20 °C for the sample in L-MNG) to prevent the protein from forming larger order species, although even at this temperature some larger species are still evident at ~8-12 S. These sedimentation coefficients are not corrected for the lower temperature so appear smaller than all other values reported for *Hi*SiaQM. The species at 4.5 S (red shading) is consistent with a monomer in DDM, $f/f_0$ = 1.1 (1σ error = 1.06–1.15) (middle panel). In contrast, a dimer species is unlikely given the unrealistic $f/f_0$ = 1.8 (1σ error = 1.68–1.82) (not shown). The larger species at 6.5 S (blue shading) is consistent with a dimer in DDM, $f/f_0$ = 1.2 (1σ error = 1.19–1.3) (right panel), whereas the monomer provides an unrealistic $f/f_0$ = 0.8 (1σ error = 0.75–0.82) (not shown). With the detergent bound to the sedimenting protein now known alongside the mass of the detergent monomers, it can be estimated that approximately 220 (monomer) and 380 (dimer) molecules of DDM are bound to the protein.

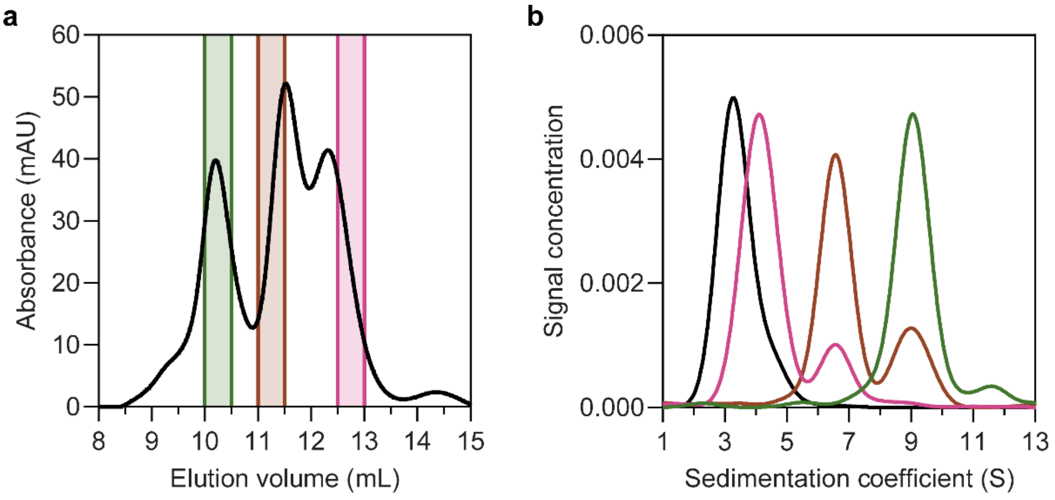


**Supplementary Figure 7 | Solubilisation of *Hi*SiaQM in nanodiscs.** **a**, The size exclusion chromatogram following nanodisc reconstitution with a 1:4:80 ratio of *Hi*SiaQM:MSP:lipid identifies the presence of multiple species. Three fractions across the elution profile were analysed with SV-AUC (shown by the coloured bars). **b**, SV-AUC analysis of nanodisc solubilised *Hi*SiaQM (pink, brown, and green) and empty nanodiscs (black) shows that later eluted fractions gave a main species consistent with empty nanodiscs (pink, 4.0 S), middle eluted fractions gave a main species consistent with a monomer incorporated (brown, 6.8 S) and earlier eluted fractions gave a larger 9.0 S main species (green). The larger species at 9.0 S is most likely a dimer of *Hi*SiaQM in a nanodisc as it has been shown to exist as a dimer in both detergent and amphipol. The dimer can just fit by physical constraints as the MSP forms ~11 nm nanodiscs and the size of the *Hi*SiaQM dimer is ~10 nm at the widest point. While physically possible, the number of reconstituted lipids would be low.

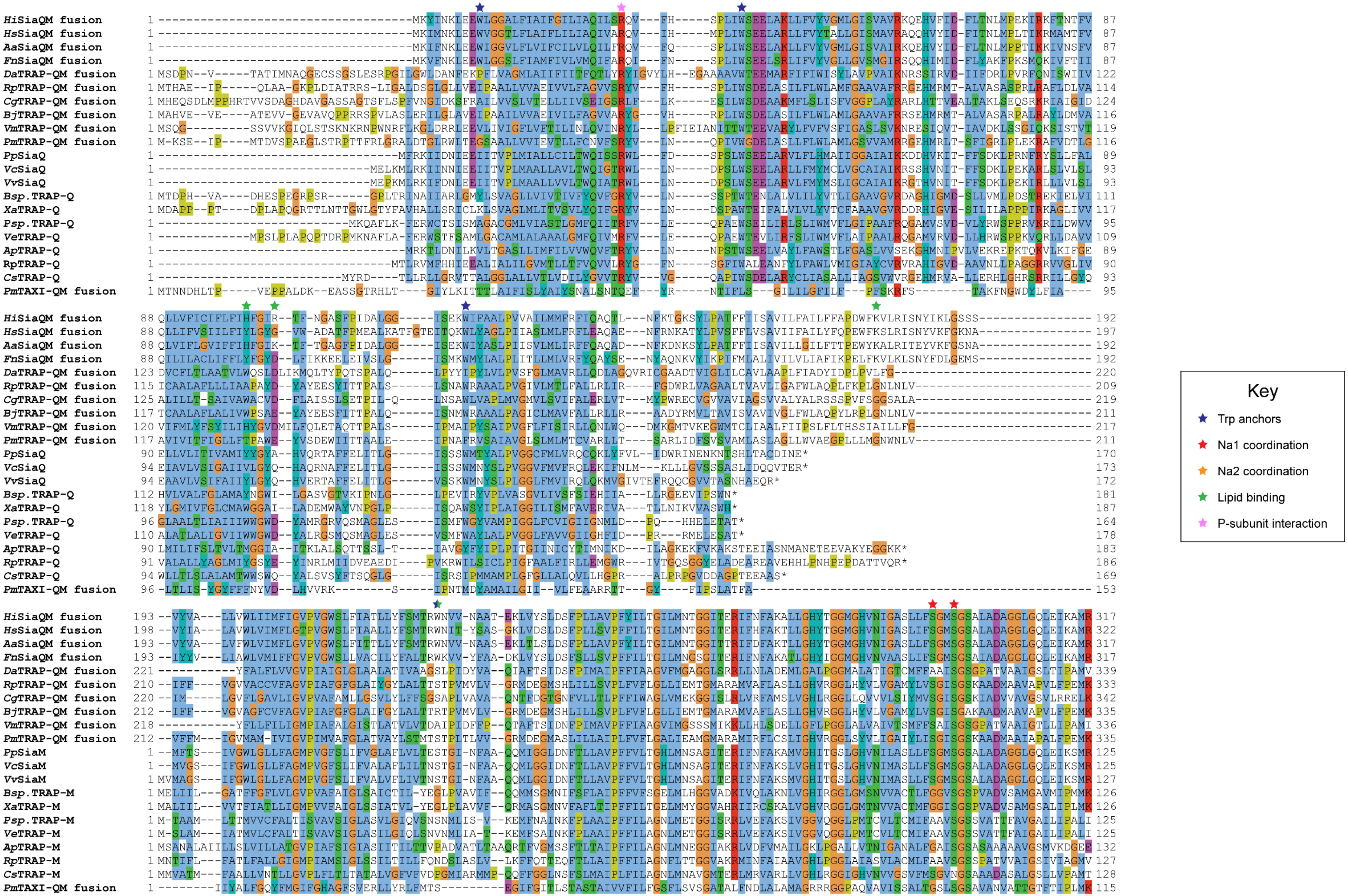
Key
Trp anchors
Na1 coordination
Na2 coordination
Lipid binding
P-subunit interaction

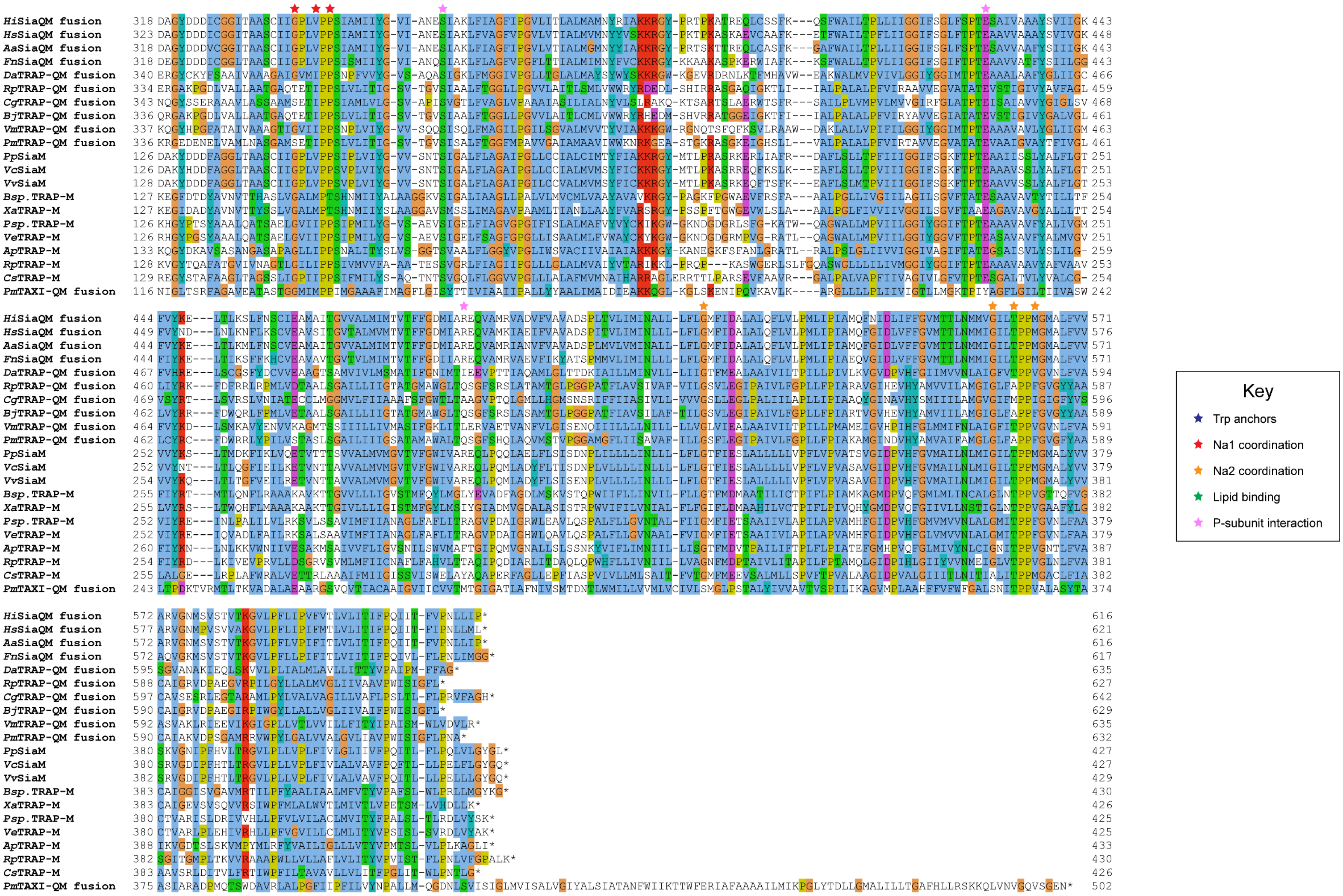
Key
Trp anchors
Na1 coordination
Na2 coordination
Lipid binding
P-subunit interaction

**Supplementary Figure 8 |** Protein sequence alignment of TRAP transporter QM-subunits. Protein sequences were obtained from the National Center for Biotechnology Information, aligned using Kalign (82) and coloured in Jalview (83) with Clustal X colouring. Anchoring tryptophan residues (blue stars), Na1 coordinating residues (red stars), Na2 coordinating residues (orange stars), predicted lipid binding residues (green stars) and expected P-subunit interacting residues (pink stars) are annotated.

SiaQM protein sequences from *Haemophilus influenzae* (*Hi*, WP_005694432), *Histophilus somni* (*Hs*, WP_249962964), *Aggregatibacter actinomycetemcomitans* (*Aa*, WP_005592763), *Fusobacterium nucleatum* (*Fn*, WP_060798477), *Photobacterium profundum* (*Pp*, WP_011218955/WP_011218954), *Vibrio vulnificus* (*Vv*, WP_001889698/WP_000233105) and *Vibrio cholerae* (*Vc*, WP_011081661/BAC97226) were aligned with TRAP-QM protein sequences from *Desulfovibrio alaskensis* (*Da*, WP_011367273), *Rhodopseudomonas palustris* (*Rp*, WP_011441572), *Caballeronia grimmiae* (*Cg*, WP_052006082), *Bradyrhizobium japonicum* (*Bj*, WP_248883330), *Virgibacillus massiliensis* (*Vm*, WP_051739028), *Pseudomonas mandelii* (*Pm*, WP_033060252), *Bradyrhizobium* sp. BTAi1 (*Bsp.*, WP_011942649/WP_011942650), *Xanthobacter autotrophicus* (*Xa*, ABS68596/ABS68595), *Polaromonas* sp. JS666 (*Psp.*, WP_011482802/WP_011482801), *Verminephrobacter eiseniae* (*Ve*, ABM59662/WP_011811650), *Anaerococcus prevotii* (*Ap*, WP_015778303/WP_015778302), *Rhodopseudomonas palustris* (*Rp*, WP_011442208/WP_011442207) and *Chromohalobacter salexigens* (*Cs*, WP_011505968/ABE58023) and the TAXI-TRAP-QM protein sequence from *Proteus mirabilis* (*Pm*, WP_004242632).

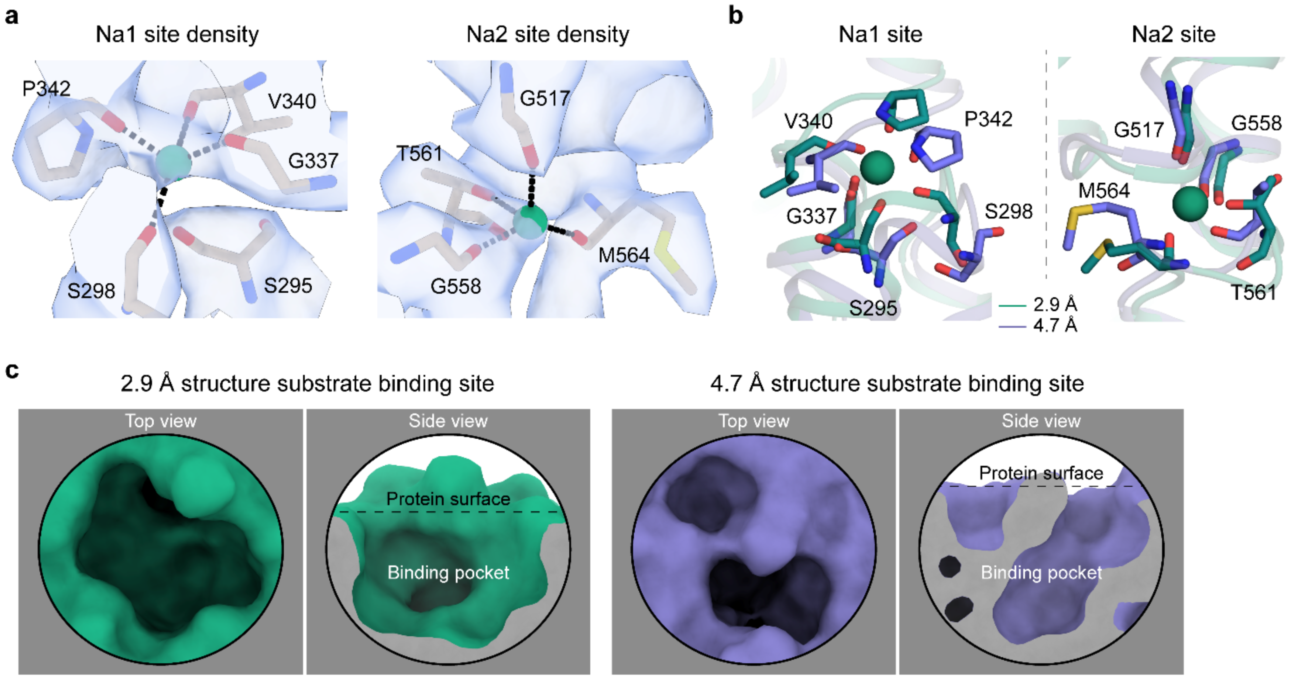


**Supplementary Figure 9 | The Na$^+$ and substrate binding sites of *Hi*SiaQM.** **a**, Closeup views of the cryo-EM density (blue) at the Na1 and Na2 sites of the antiparallel *Hi*SiaQM dimer. Density is present for both Na$^+$ ions (green spheres). Na$^+$ binding residues are shown as sticks and Na$^+$ coordination as black dashes. The density map was automatically sharpened using Phenix (84) and the radii of the Na$^+$ ions have been reduced to show the density more clearly. **b**, A comparison of the Na$^+$ binding sites of *Hi*SiaQM from the 2.9 Å structure (green) and the 4.7 Å structure (purple). The Na$^+$ ions of the 2.9 Å structure are shown as green spheres and are well positioned to coordinate the identified binding residues in the 'clamshell' loops. The positions of these loops are different between the two structures. **c**, A comparison of the putative Neu5Ac binding sites of *Hi*SiaQM from the 2.9 Å structure (green) and the 4.7 Å structure (purple). The higher resolution structure shows that a defined binding pocket exists at the centre of the transport domain.

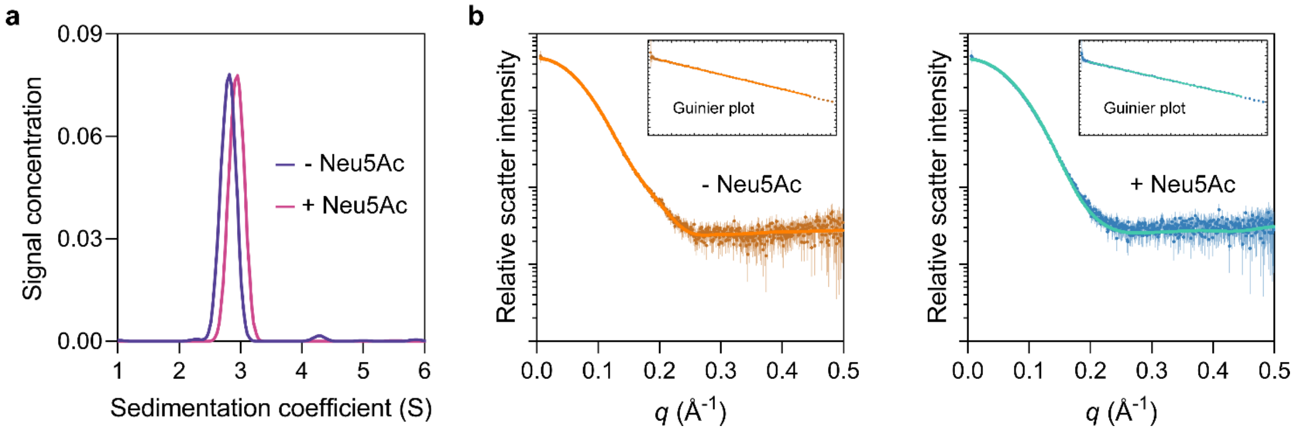


**Supplementary Figure 10 | *Hi*SiaP is monomeric.** **a**, Analytical ultracentrifugation experiments of *Hi*SiaP (expressed in the periplasm) without Neu5Ac or with 5 mM Neu5Ac resulted in sedimentation coefficients of ~2.8 S without Neu5Ac (blue) and ~3.0 S with Neu5Ac (pink), consistent with a monomeric oligomeric state. The samples had calculated masses of 39 and 35 kDa, indicating that *Hi*SiaP exists as a monomer in both a ligand bound and unbound state. **b**, SAXS analysis of *Hi*SiaP without Neu5Ac (left, brown) and with 5 mM Neu5Ac (right, blue) at a considerably higher protein concentration than analytical ultracentrifugation (360 µM) resulted in a radius of gyration ($R_g$) of 20.9 Å and a maximum particle dimension ($D_{max}$) of 64 Å without Neu5Ac and 20 Å and 63 Å with 5 mM Neu5Ac. CRYSOL was used to fit the predicted scatter from monomeric crystal structures to the observed scatter (74). Fitting *Hi*SiaP without Neu5Ac (PDB: 2CEY) (left, orange) results in $X^2$ = 0.32. Fitting *Hi*SiaP with Neu5Ac (PDB: 3B50) (right, teal) results in $X^2$ = 0.66. Fitting the incorrect structures results in $X^2$ values of 4.91 (without Neu5Ac data to 3B50) and 2.36 (with Neu5Ac data to 2CEY). These SAXS analyses of *Hi*SiaP are all consistent with the sizes of the monomers from crystallography (23,24).

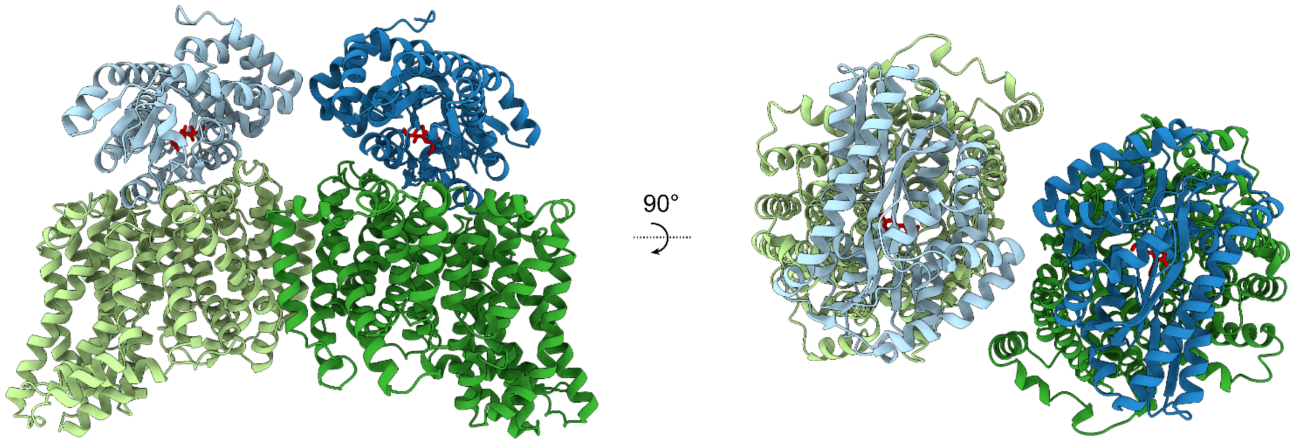


**Supplementary Figure 11 | Modelling the *Hi*SiaPQM complex.** AlphaFold2 was used to model the complex of two *Hi*SiaP monomers (PDB: 3B50) bound to the parallel *Hi*SiaQM dimer from this work (PDB: 8THI). The *Hi*SiaQM monomers are shown in two shades of green and the *Hi*SiaP monomers (ligand bound, closed form) are shown in two shades of blue. Neu5Ac bound to *Hi*SiaP (red sticks) is positioned directly over top of the interface between the scaffold and transport domains of each *Hi*SiaQM monomer. The predicted binding mode of the complex allows for the formation of two complete tripartite *Hi*SiaPQM systems without significant steric clashes.

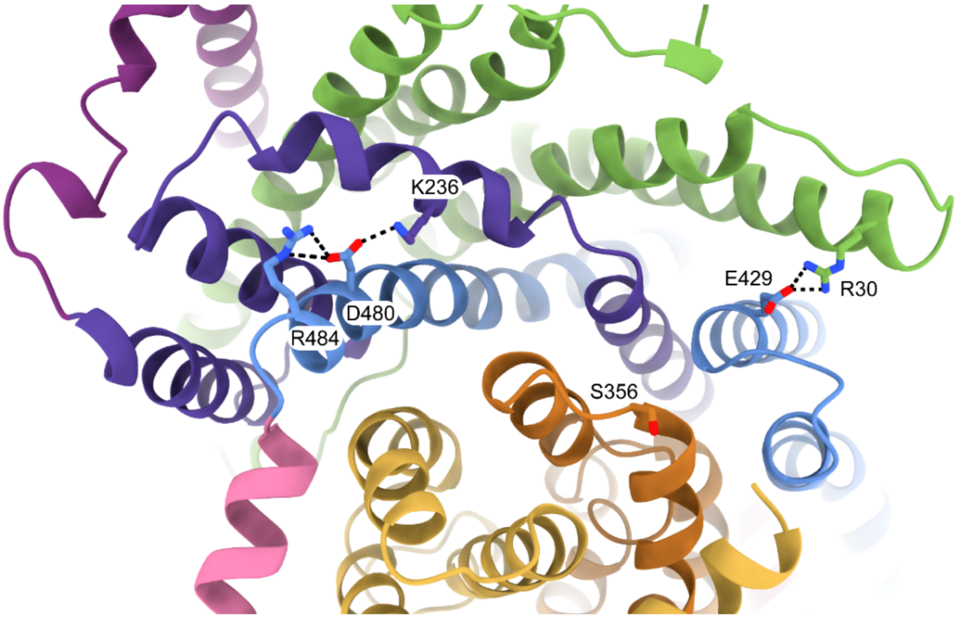


**Supplementary Figure 12 | Residues involved in scaffold stabilisation and the interaction with SiaP.** The resolution of the *Hi*SiaQM structure (coloured as in **Figure 2c**) shows the positions of R30, S356, E429 and R484 as viewed from the periplasm. R30 (SiaQ, helix 1) forms a salt bridge with E429 (SiaM, helix 8) to connect two helices of the scaffold and SiaQ and SiaM. S356 is located on the loop between helices 5b and 6 of SiaM in the transport domain. R484 (SiaM, helix 9a) forms a salt bridge with D480 (SiaM, helix 9a), which interacts with K236 (SiaM, helix 3a) to connect two helices of the scaffold.

**Supplementary Movie 1 |** Three-dimensional variability analysis of our *Hi*SiaQM reconstructions shows only very subtle motion at the dimer interface, and does not show any global elevator motions in the final reconstruction.
